# BLA^KOR^ inputs to the BNST regulate social stress-escalated alcohol consumption

**DOI:** 10.1101/2024.11.07.622470

**Authors:** Franciely Paliarin, Chelsea Duplantis, Evan Doré, Samhita Basavanhalli, Emma Weiser, Tameka W. Jones, Rajani Maiya

**Affiliations:** Department of Physiology, Louisiana State University Health Sciences Center, New Orleans, LA, 70112

**Keywords:** *Oprk1-Cre*, NorBNI, basolateral amygdala, kappa opioid receptor, chemogenetics, social defeat stress, intermittent access

## Abstract

**Background:** Aversive social experiences can lead to escalated drug consumption and increase the risk of relapse to drug seeking. Individuals who consume alcohol to alleviate the effects of social stress are more likely to develop an alcohol use disorder (AUD). Repeated social defeat stress (SDS) enhances the rewarding and reinforcing effects of alcohol. However, the neural mechanisms that underlie social stress-escalated alcohol drinking are not well understood. Here we explored the role of the dynorphin/kappa opioid receptor (Dyn/KOR) system in regulating social stress-escalated alcohol consumption.

**Methods:** Male and female mice were subjected to repeated SDS for 10 days following which they were left undisturbed in their home cages. They were then subject to intermittent access (IA) two-bottle choice alcohol consumption procedure. The effects of systemic and BNST-specific KOR antagonism using the selective KOR antagonist NorBNI on stress-escalated drinking were evaluated. Using chemogenetic approaches in *Oprk1-Cre* mice, we examined the role of KOR expressing cells in the basolateral amygdala (BLA^KORs^) and BLA^KOR^-BNST pathway in social stress-escalated alcohol consumption.

**Results:** Repeated SDS increased alcohol consumption and preference in both males and females. Systemic KOR antagonism attenuated SDS-escalated alcohol consumption in both males and females. BNST -specific KOR antagonism also attenuated stress-escalated drinking in males. Finally, selective chemogenetic activation of BLA^KORs^ and BKA^KOR^-BNST pathway attenuated social stress-escalated alcohol consumption in both sexes.

**Conclusion:** Our results suggest a significant role for BLA^KOR^ projections to the BNST in regulating social stress-escalated alcohol consumption. Our results provide further evidence that the Dyn/KOR system maybe a viable target for medications development to tareat comorbid stress and AUD.

## Introduction

Social stress is pervasive in humans and its severity is correlated with a wide variety of neuropsychiatric conditions including mood and substance use disorders (1⍰3). Exposure to repeated social stress is associated with escalated drug intake and increased risk of relapse to drug seeking for a variety of drugs including cocaine and alcohol (2, 4). The link between social stress and alcohol misuse is well established. Individuals that consume alcohol to alleviate social anxiety and stress are more likely to meet the Diagnostic and Statistical Manual (DSM-V) criteria for an alcohol use disorder (AUD) (3). Further, exposure to moderate or severe psychosocial stressors is associated with increased likelihood of developing a sustained pattern of uncontrollable drinking in those who drink as a coping mechanism (3, 5⍰7). The effects of social stress are evaluated in the clinic through the Trier social stress task (TSST) which reliably increases cortisol levels and heart rate in response to social examination and performance stress. TSST results in increased craving and responding to alcohol in patients suffering from AUD suggesting that these individuals are more likely to misuse alcohol in order to cope with negative affective situations (8⍰10). The high comorbidity between stress and AUD (11⍰15) underscores the importance of studying molecular mechanism underlying the effects of stress on alcohol use.

There are several types of psychosocial stressors including social exclusion, social defeats stress, and social isolation stress (4). Social defeat stress (SDS) is modeled in rodents using resident-intruder paradigms (2, 16, 17). Though the mechanics of SDS may vary, the physiological and behavioral effects of SDS are remarkably similar across species (18). Repeated brief episodes of SDS reliably leads to long-lasting increases in alcohol consumption and preference in both male and female mice (2, 19, 20). Several studies have examined the neural mechanisms by which SDS leads to escalated drug use. SDS can sensitize brain dopamine (DA) systems thereby enhancing vulnerability to drug use (2). SDS activates several neural and endocrine processes (2, 4, 21⍰23) and mobilizes peptidergic systems (22, 24, 25) including CRF (21, 24), and Orexin (24).

The opioid peptide Dynorphin (Dyn) and its cognate receptor the Kappa opioid receptor (KOR) have been implicated in neural processes by which stressful situations precipitate relapse and reinstatement to drug seeking (26⍰31). KOR antagonists prevent reinstatement to drug seeking by a variety of stressors and ameliorate depressive-like states induced by drug withdrawal (27, 32⍰38). Consistent with these findings, the short acting KOR antagonist Aticparant is currently in clinical trials for treating major depressive disorder (39). KOR agonists are aversive and can mimic stress and lead to increased anxiety-like behaviors and drug seeking (40⍰42).

However, the role of Dyn/KOR system in regulating social stress-escalated alcohol consumption has not been evaluated. KOR is a member of the seven transmembrane G protein-coupled receptor superfamily coupled to the G_i/o_ subunit. Dyn-mediated activation of KOR leads to inhibition of neuronal activity by reducing calcium currents, activating potassium channels, and decreasing cyclic AMP (cAMP) (26, 43, 44). The Dyn/KOR system is expressed in several brain regions including those implicated in the behavioral response to stress such as the central (CeA), basolateral amygdala (BLA), and the bed nucleus of the stria terminalis (BNST). KOR’s are typically expressed on presynaptic terminals where they are poised to regulate the release of neurotransmitters and neuromodulators such as glutamate, DA, and serotonin (5-HT) (26, 27, 45). Dyn is a stress responsive neuropeptide and is released in response to a variety of stressors (27). Dyn-mediated activation of KORs and subsequent β-arrestin dependent phosphorylation of p38 MAPK (33) mediates the aversive and dysphoric components of stress. Activation of KORs can also be anxiogenic. Specifically, activation of KORs located on BLA terminals in the BNST gate anxiety-like behaviors (44). Upregulation of Dyn/KORs are also thought to be part of opponent processes activated during drug withdrawal that contribute to escalated drug intake in dependent animals

In this study, we evaluated the role of Dyn/KOR system in regulating SDS-escalated alcohol consumption in both male and female mice. Our results indicate a role for Dyn modulation of BLA^KOR^ inputs to the BNST in regulating SDS-escalated alcohol consumption in both male and female mice.

## Methods

### Animals

7-8 weeks old male and female C57BL/6J mice were purchased from The Jackson Laboratory (Bar Harbor, ME). The generation and characterization of *Oprk1-Cre* mice is described (46). *Oprk1-Cre* mice were backcrossed to C57BL/6J mice for at least 3-4 generations before being used in the study. Mice were provided access to food and water *ad libitum* and individually housed. Mice were maintained on Tl2019S (Teklad, Innotiv) chow throughout the drinking study or switched to LD5053 chow from Research Diets (New Brunswick, NJ) at least 48 hours prior to the start of the drinking study. Mice maintained on LD5053 chow showed higher levels of alcohol intake compared to those maintained on TL2019S diet (47). Mice were allowed to habituate to the reverse light/dark cycle (on 10:00 A.M., off 10:00 P.M.) for 1-week prior to initiation of experiment. During the habituation period, mice were individually housed in double grommet cages equipped with two bottles provided with sipper tubes containing water. Animals were weighed once per week at the beginning of the week.

### Drugs

Norbinaltrophimine (NorBNI) was purchased from Tocris Pharmaceuticals (Minneapolis, MN) and dissolved in sterile saline and administered at 10 mg/kg (10 ml/kg) intraperitoneally or 2.5 μg/side into the BNST. Clozapine-N-Oxide (CNO, freebase) was purchased from HelloBio (Princeton, NJ). CNO was dissolved in saline containing 0.5% Dimethyl Sulfoxide (DMSO). CNO was administered at a dose 3 mg/kg (10 ml/kg) intraperitoneally and 3 μM into the BNST.

### Social defeat stress

Mice were subject to repeated SDS as described (17). In this protocol, eight-week-old individually housed male (C57BL/6J or *Oprk1-Cre*) mice served as intruders. Aggressive Swiss Webster (CFW) males housed with tubally ligated or ovariectomized CFW females served as residents. Intruders were randomly assigned to non-defeated control or social defeat groups. Each defeat episode lasted for a maximum of 15 minutes and comprised of three phases: 1) pre-defeat threat period that lasts for 3 minutes, 2) SDS episode which lasts for a maximum of 5 minutes, and 3) post-defeat threat period that lasts for 5 minutes. During the pre-defeat threat period, CFW female was temporarily removed from CFW male’s home cage, and the intruder mouse was introduced in a perforated protective plexiglass box. The protective box around the intruder was removed to initiate social defeat. Intruders were directly exposed to the resident until they received 30 bites/attacks, or a total of 5 minutes had elapsed. Following defeat, intruders were placed back in the protective plexiglass box while remaining in the resident’s home cage for the post-defeat threat period for 5 minutes after which they were returned to their home cage. Mice were subject to SDS once every day for ten days and encountered a new CFW male for each defeat session. Following SDS, mice were left undisturbed in their home cages for 10 days after which they were subject to intermittent access (IA) alcohol consumption. Social defeat stress in females is notoriously difficult to implement as under most conditions neither male nor female residents attack female intruders. We used a method recently developed in Dr. Miczek’s lab where resident CFW females were housed with castrated CFW males for 1-week prior to the start of defeat (19). CFW females housed with castrated males become dominant/aggressive and will reliably attack intruders introduced into their home cage. The male CFW mouse was temporarily removed, and a female intruder mouse was introduced into the CFW female’s home cage. Defeat sessions were conducted exactly as outlined for males with the exception that defeats were stopped once female intruders received 20 attacks. This procedure results in reliable and robust social defeat in females (19).

### Intermittent access (IA) two bottle choice alcohol consumption

Male and female mice were subjected to IA alcohol consumption procedure as previously described (48). Mice were maintained on TL2019S diet and switched to LD5053 at least 48 hours before the start of the drinking study. Initial cohorts of mice were maintained on TL2019S throughout the IA alcohol procedure. Animals were given two bottles from which to drink, one containing water and the other containing15% ethanol (V/V) (PHARMCO-AAPER, Brookfield, CT) solution for 24 hours every other day. On the off-days, mice had access to two bottles of water. Alcohol and water bottles were weighed prior to placing in the cage and were weighed again 24-hours later. Spillage was controlled for by measuring volume lost from alcohol and water bottles placed on an empty cage. The position of the water and alcohol bottles were switched for every drinking session. Control water drinking mice were provided with two bottles of water. Mice were weighed once per week.

### Stereotaxic surgeries

AAVs (150-200 nl) encoding either Cre-dependent mCherry (AAV8-hSyn-DIO-mCherry) or hM3DQ (AAV8-hSyn-DIO-hM3DQ, from Addgene, Waterstown, MA) as described previously (46) were infused bilaterally into the BLA of *Oprk1-Cre* mice using a Nanoject III (Drummond Scientific) injector at the rate of 25 nl/minute. The coordinates for BLA were A/P −1.6, M/L ±3.25, D/V −4.4 from skull. The titer of the virus injected was approximately 1 x 10^12^ vg/ml. A separate group of mice were implanted with stainless steel guide cannula (Plastics one, Roanoke, VA and RWD Life Science, Sugarland, TX) aimed bilaterally at the BNST (AP +0.3, ML ± 0.1, DV −4.3 from skull). Cannulas were affixed to the skull with dental acrylic (H00325, Coltene, Cuyahoga Falls, OH). Drugs were infused using an injector whose tip protruded 0.1 mm past the guide cannula. All mice were allowed to recover for 1-2 weeks prior to behavioral testing.

### Statistical analysis

All data are presented as the mean +/− SEM. Statistical analysis was assessed using unpaired t-tests and two- or three-way repeated measures (RM) ANOVA. Posthoc Sidak tests were performed when appropriate significant main effects or interactions were detected. For chemogenetics and BNST-specific KOR antagonism experiments, only data from subjects with verified correct placements of viruses or microinjectors were used for analysis.

## Results

### Repeated SDS leads to robust increases in alcohol consumption in both sexes

We first determined if repeated SDS reliably led to escalated alcohol consumption in male and female C57BL/6J mice. Male and female mice were subject to SDS for 10 days as outlined in methods following which mice were left undisturbed in their home cages for 10 days. They were then exposed to IA alcohol procedure for 3 weeks. Two-way repeated measures (RM) ANOVA of average weekly alcohol consumption in males (**Fig. 1A**) revealed a significant main effect of stress [F_Stress_ (1,62) = 22.09, P<0.001], session [F_Session_ (2, 124) = 8.776, P = 0.0003], and a marginally significant stress x session interaction [F_Stress x Session_ (2, 124) = 2.801, P = 0.065]. Posthoc Sidak test revealed that male mice consumed significantly more alcohol per 24-hour session on all 3 weeks compared to unstressed controls (***, P<0.001; **, P<0.01; *, P<0.05). Two-way RM ANOVA of alcohol preference (**Fig. 1B**) also revealed a significant main effect of stress [F_Stress_ (1,62) = 26.77, P<0.001], drinking session [F_Session_ (2,124) = 23.13, P<0.001], and stress x session interaction [F_Stress_ (2, 124) = 3.303, P<0.001]. Posthoc Sidak test revealed that stressed male mice showed enhanced preference for alcohol across all 3 weeks (***, P<0.001; **; P<0.01; *, P<0.05). Two-way RM ANOVA analysis of alcohol consumption in females (**Fig. 1C**) also showed a significant main effect of stress [F_Stress_ (1, 45) = 7.132, P = 0.0105] and session [F_Session_ (2, 90) = 16.48, P <0.0001]. Posthoc Sidak’s test showed that stressed females consumed significantly higher amounts of alcohol on week 2 compared to unstressed controls (*, P<0.05). Two-way RM ANOVA analysis of alcohol preference (**Fig. 1D**) in females revealed a significant main effect of stress [F_Stress_ (1,45) = 9.456, P = 0.0036] and session [F_Session_ (2, 90) = 4.715, P = 0.0113]. Sidak’s post-hoc test revealed that female mice displayed significantly higher preference for alcohol during the first two weeks of the IA procedure (**, P<0.01; *, P<0.05). To more directly compare males and females, we represented the data as % increase in drinking in stressed mice compared to controls in males and females across 3 weeks of IA alcohol procedure (**Fig. S1**). We did not find significant differences in the magnitude of stress-escalated drinking between males and females across sessions suggesting that repeated SDS increased alcohol consumption to a similar extent in both males and females. In summary, these results indicate that repeated SDS led to robust and persistent increases in alcohol consumption and preference in both sexes.

**Figure 1:**
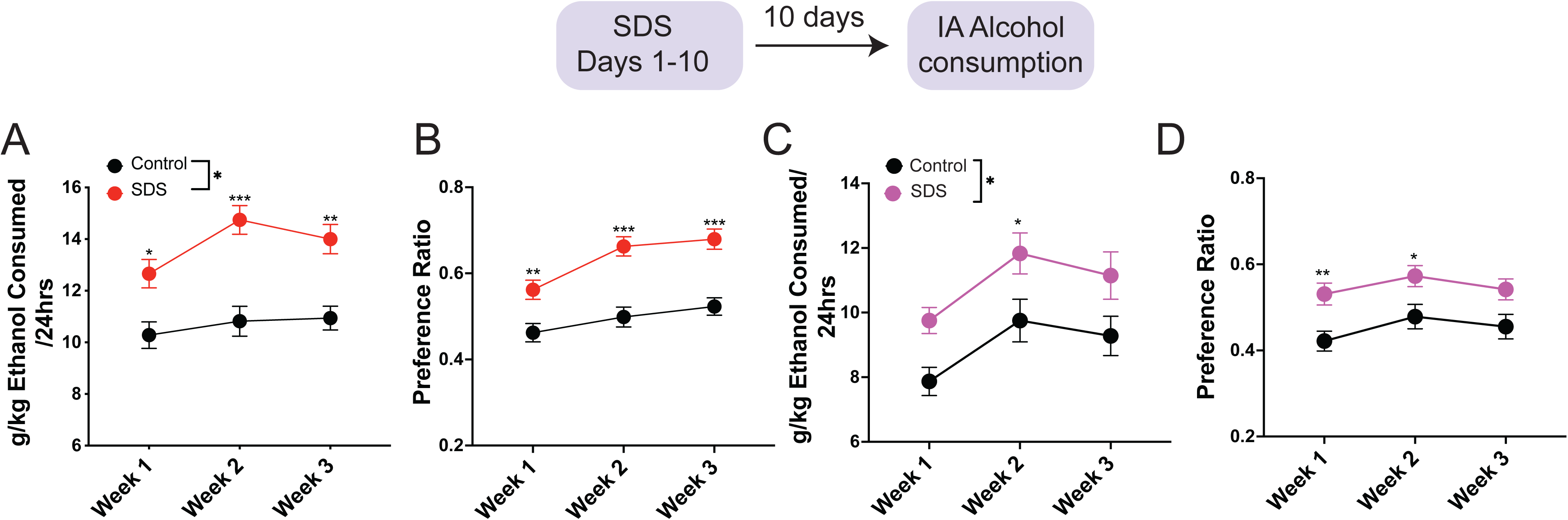
Repeated social defeat stress results in long lasting increases in alcohol consumption and preference in **males** (**A**&**B**) and **females** (**C**&**D**). Average alcohol consumption and preference per session per week are shown. Social defeat stress significantly increased **A)** alcohol consumption (*, p<0.05, **, P<0.01, Sidak’s post-test) and alcohol preference **(B)** in males. (**, p<0.01, ***. P<0.0001, Sidak’s post-test). N =26-35 mice /group. Repeated social defeat stress alco significantly increased **(C)** alcohol consumption *, P<0.05, Sidak’s post-test and preference **(D)** in females. **, P<0.01; P<0.05, Sidak’s post-test, N = 23-24 mice/group for females.

### Systemic NorBNI injections attenuate social stress-escalated alcohol consumption in both sexes

We next determined the role of Dyn/KOR interactions in social stress-escalated alcohol consumption. We first examined whether systemic administration of Norbinaltorphimine (NorBNI), a selective KOR antagonist attenuated stress-escalated alcohol consumption in both sexes. Mice were subject to SDS followed by IA alcohol procedure for 3 weeks (**Fig. 2A**). Mice were then administered vehicle or NorBNI (10 mg/kg, i.p.) systemically 24-hours prior to the alcohol drinking session in a within subjects’ design. Alcohol and water consumption was measured 3 hours (males) and 4 hours (females) after alcohol bottles were introduced on the first (S1) and second (S2) alcohol drinking session post vehicle or NorBNI administration (**Fig. 2B-E**). Since NorBNI-mediated inhbition of KOR function is long lasting, we were unable to counterbalance vehicle and drug injections in this study. Two-way RM ANOVA analysis of alcohol consumption (**Fig. 2B**) revealed a significant stress x drug interaction [F_Stress x Drug_ (2, 48) = 6.320, P = 0.0037]. Posthoc Sidak test revealed that NorBNI injections significantly reduced alcohol consumption in stressed mice compared to vehicle injections across both alcohol drinking sessions (S1 and S2; **, P<0.01; *, P<0.05). NorBNI did not affect alcohol consumption in unstressed controls compared to vehicle injections. NorBNI did not impact water consumption across sessions in both stressed and control mice (**Fig. 2C**).

**Figure 2:**
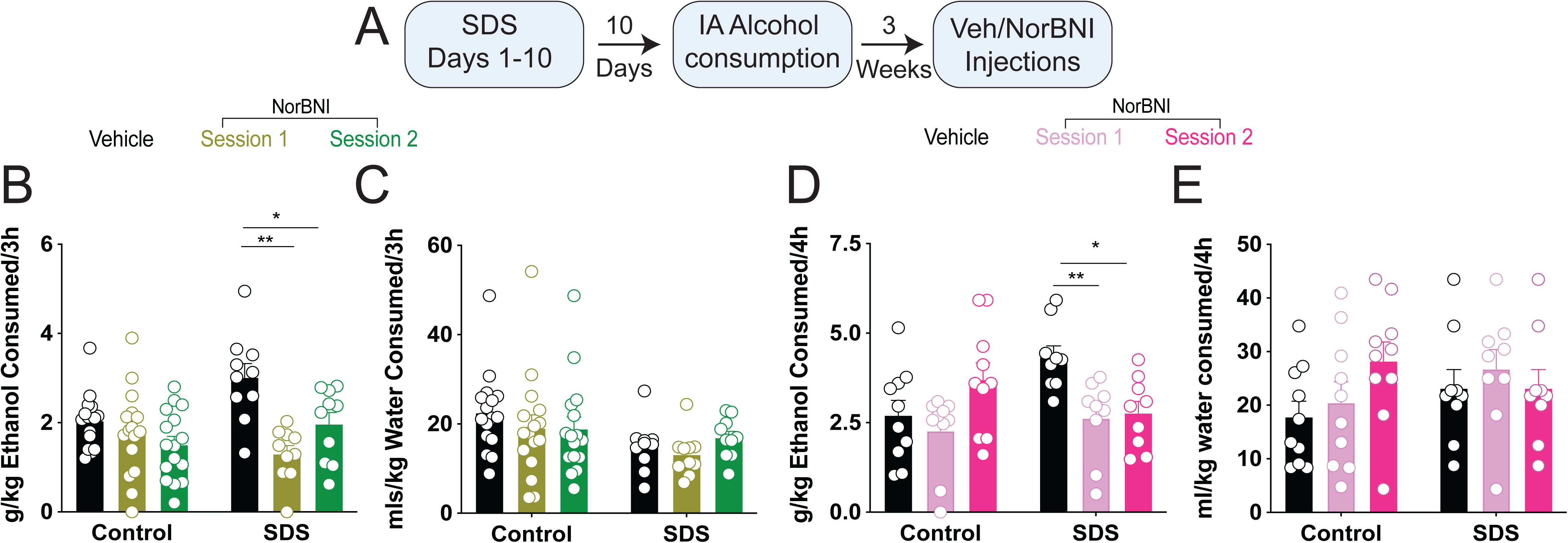
Systemic administration of a selective KOR antagonist NorBNI reduced alcohol consumption in stressed **males** (**B** and **C**) and **females** (**D** and **E**) but not unstressed controls. **A**) Experimental timeline is shown. **B)** Alcohol consumption is shown for two sessions after NorBNI administration in males. Sidak’s post-test revealed that NorBNI significantly reduced alcohol consumption across both sessions. **, p<0.01, *, p<0.05, compared to vehicle injected mice. N = 10-15mice/group. **C)** Water consumption was not affected in males. **D**) NorBNI reduced alcohol consumption in stressed females but not in unstressed controls. **, p<0.01, *, p<0.05, Sidak post-test. N = 9-10 mice /group. **E**) Water consumption was not affected in females.

Similar to males, NorBNI also significantly reduced alcohol intake in stressed (**Fig. 2D**) but not in unstressed females. Two-way RM ANOVA of alcohol consumption indicated a significant stress x drug interaction [F_Stress x Drug_ (2, 34) = 3.287, P = 0.0495], and a significant main effect of stress [F_Stress_ (2, 34) = 8.779, P = 0.0008]. Posthoc Sidak analysis revealed that NorBNI significantly reduced alcohol consumption in first session (S1) post drug injection in stressed mice but not in unstressed controls (**, p<0.01; *, p<0.05). NorBNI did not affect water consumption in either stressed or control females (**Fig. 2E**). We also found similar trends in effects of NorBNI on alcohol consumption at the 24h timepoint (data not shown). In summary, these results suggest that NorBNI treatment selectively attenuates alcohol consumption in stressed males and females but not in unstressed controls thereby implicating a role for KORs in regulating stress-escalated drinking.

### Chemogenetic activation of BLA^KOR^ neurons attenuates SDS-escalated alcohol consumption in males

We next asked where in the brain KORs may be functioning to regulate social stress-escalated alcohol drinking. To address this question, we used the newly generated and characterized *Oprk1-Cre* mouse (46) which expresses *Cre* recombinase in *Oprk1* cells throughout the brain with high fidelity. We first focused on the BLA as it plays an important role in mediating the behavioral effects of social stress (49, 50), regulates alcohol consumption (51, 52), and expresses high levels of KORs (51). We hypothesized that stress-induced increases in Dyn would inhibit KOR-expressing neurons in the BLA. We therefore tested the prediction that chemogenetic activation of BLA^KOR^ neurons would counter this inhibition and attenuate stress-escalated drinking. AAVs encoding Cre-dependent mCherry or hM3DQ were injected into the BLA of *Oprk1-Cre* mice (**Fig. 3A**). There were 4 experimental groups: 1) Control -mCherry, 2) SDS – mCherry, 3) Control - hM3DQ, and 4) SDS - hM3DQ. Mice were subjected to SDS followed by IA alcohol consumption for 3 weeks following which all mice were administered vehicle and CNO injections (i.p.) in a within-subjects’ Latin-square design 30 minutes prior to the start of the drinking session (**Fig. 3B**). Three-way RM ANOVA revealed a significant stress x virus x drug interaction [F_stress x virus x drug_ (1,27) = 7.428, P = 0.011]. Posthoc Sidak test revealed that CNO (3 mg/kg i.p) significantly reduced alcohol consumption 3-hours post injection in stressed hM3DQ-injected mice compared to vehicle controls (****, P<0.001, **Fig. 3A**). CNO administration also significantly reduced alcohol consumption in control unstressed hM3DQ-injected mice compared to vehicle controls (*, P<0.05, **Fig. 3C**). This result is consistent with literature suggesting a role for KORs in regulating basal alcohol consumption (41, 53). To determine if there was a difference in magnitude of CNO-mediated reduction in alcohol intake between control and stress groups, we represented the data as difference in drinking between vehicle and CNO-injected mice for each group (**Fig. 3E**). We found that the magnitude of reduction in drinking was greater in stressed mice compared to unstressed controls suggesting greater engagement of KORs in stressed mice compared to unstressed controls. Two-way ANOVA of difference in drinking between vehicle and CNO injected groups showed a significant stress x virus interaction [F_stress X virus_ (1,27) = 5.865, P =0.0224]. Posthoc Sidak tests revealed that CNO administration significantly reduced alcohol drinking in stressed hM3DQ-injected mice compared to control hM3DQ-injected mice (*, P<0.05). Alcohol consumption was also significantly reduced in stressed hM3DQ-injected mice compared to stressed mCherry-injected mice (***, P< 0.001). Water consumption was not affected in any of the groups (**Fig. 3D**). These results implicate a role of BLA^KOR^ neurons in regulating both stress-escalated and basal alcohol consumption.

**Figure 3:**
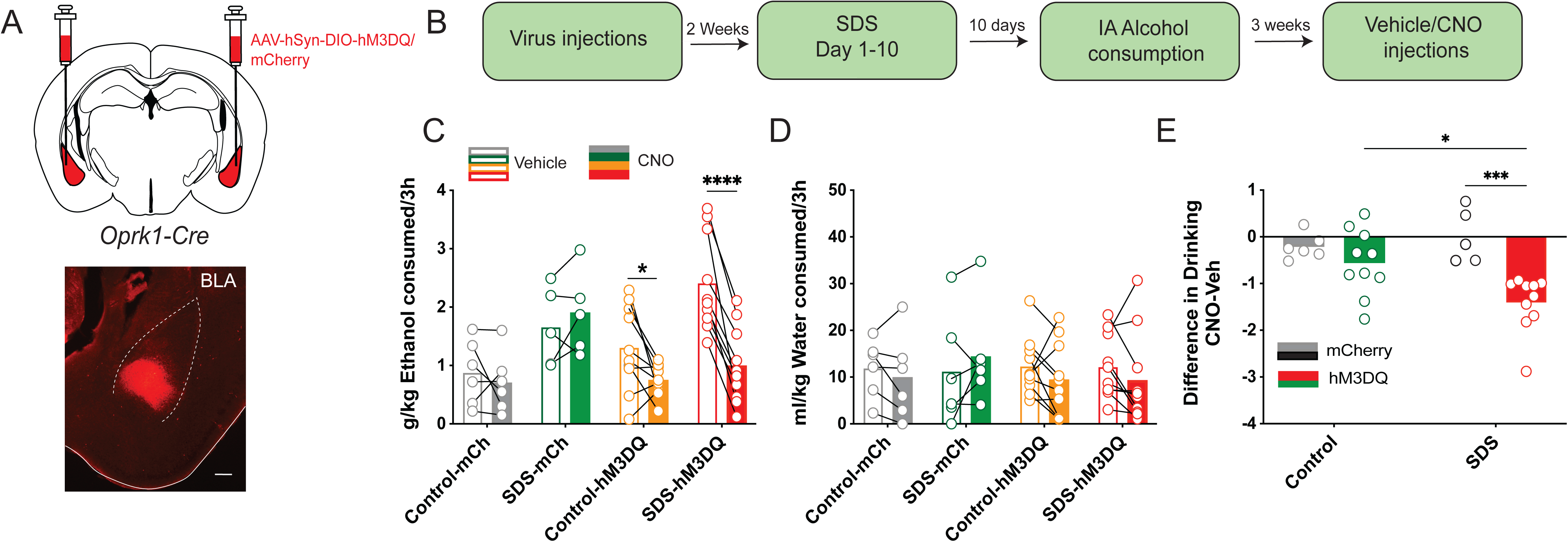
Chemogenetic activation of BLA^KOR^ cells significantly attenuated alcohol consumption in hM3DQ-injected stressed mice and unstressed control males. However, the magnitude of reduction in drinking was greater in stressed hM3DQ-injected mice. **A)** Schematic of virus targeting to BLA in *Oprk1-Cre* mice. Scalebar = 100 μm. **B)** Timeline of SDS and IA alcohol procedure and CNO injections is shown. **C**) CNO administration significantly reduced alcohol consumption. Posthoc Sidak’s test revealed that CNO administration significantly reduced alcohol consumption in both stressed mice and in unstressed controls. *, P<0.05, ***, p<0.001. **D**) Water consumption was not affected in any of the groups. **E)** Change in alcohol consumption from respective vehicle alcohol intake across treatment groups is shown. Posthoc Sidak’s test revealed that CNO administration significantly attenuated alcohol consumption in stressed hM3DQ-injected mice compared to stressed mCherry injected mice. Additionally, CNO injections reduced alcohol intake to a higher degree in stressed hM3DQ-injected mice compared to control hM3DQ-injected mice. *, p<0.05, ***, P <0.001, N = 5-6 mice/group for mCherry and 10 mice/group for hM3DQ.

### BNST-specific KOR antagonism attenuates SDS-escalated alcohol consumption in males

We next determined downstream projection targets of BLA^KOR^ neurons that might regulate social stress-escalated alcohol consumption. BLA^KOR^ neurons project robustly to the BNST which is known to play an important role in regulating stress responsivity and alcohol intake **(54, and Fig. 5A**). Further, BLA glutamatergic inputs to the BNST are modulated by Dyn and gate anxiety-like behaviors (44). Consistent with published literature, BLA^KOR^ neurons project primarily to the anterodorsal BNST while completely avoiding the oval nucleus (44, 46, 55).

We next determined the effects of BNST-specific KOR antagonism on social stress-escalated alcohol consumption. We subjected male C57BL/6J mice to SDS followed by IA alcohol consumption. After 3 weeks of IA alcohol consumption, mice were cannulated with guide cannulas aimed over the BNST (**Fig. 4A**). Mice were allowed to recover for 1 week following which they were subjected to IA alcohol consumption for 1 week (**Fig. 4B**). Mice were bilaterally infused with vehicle and NorBNI (2.5 μg/side) in a within-subjects design 24-hours prior to the drinking session. Two-way RM ANOVA of alcohol consumption revealed a significant stress x drug interaction [F_Stress X Drug_ (1,36) = 4.460, P =0.0417] (**Fig. 4C**). Posthoc Sidak test indicated that NorBNI significantly reduced alcohol consumption in stressed mice (***, P<0.001) but not in unstressed controls. We found a small compensatory increase in water consumption in both stressed and control mice (**Fig. 4D**). Two-way RM ANOVA analysis of water consumption revealed a significant main effect of drug [F_Drug_ (1, 36) = 29.56, P < 0.0001]. Posthoc SIdak test indicated that NorBNI significantly reduced water consumption in both control and stressed mice compared to vehicle (***, P<0.001). However, total fluid intake was unaffected. These results suggest that KORs in the BNST regulate stress-escalated alcohol consumption (**Fig. 4E**).

**Figure 4:**
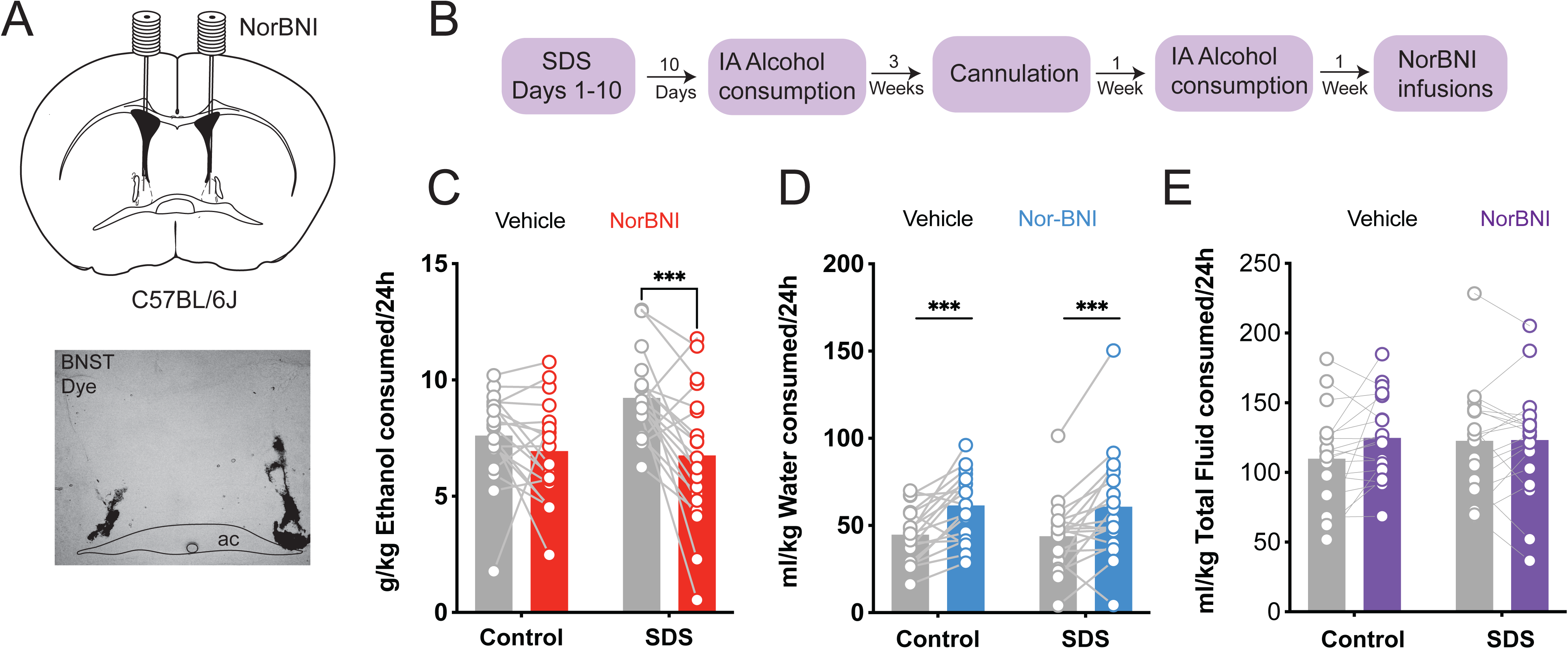
KOR antagonism in the BNST attenuated social stress-escalated alcohol consumption in males. **A)** Schematic of NorBNI infusions and representative image of cannula placement in the BNST. Scalebar = 100 μm. **B)** Experimental timeline of SDS and NorBNI infusions into the BNST. **C**) Infusion of NorBNI into the BNST reduced alcohol consumption in stressed mice but not in unstressed controls. ***, P<0.001, Sidak’s post-test, N = 19/group. **D**) There was a small but significant increase in water consumption after NorBNI-infusion in both control and stressed mice. ***, P<0.00, Sidak’s post-test, N = 19 mice/group. **C**) Total fluid consumption was not affected.

### Pathway specific activation of BLA^KOR^ inputs to the BNST attenuated SDS-escalated alcohol consumption in both sexes

We next determined whether selective chemogenetic activation of BLA^KOR^-BNST pathway attenuates social stress-escalated alcohol drinking. We injected male and female *Oprk1-Cre* mice with AAVs encoding Cre-dependent hM3DQ. Representative schematic of virus injection and cannula placement are shown in top panel **Fig. 5A**. Bottom panels show viral expression in the BLA of *Oprk1-Cre* mouse and BLA^KOR^ projections to BNST. Mice were allowed to recover and subjected to SDS, or control conditions followed by IA alcohol consumption for 3 weeks to allow the escalated drinking phenotype to emerge (see experimental timeline in **Fig. 5B**). Mice were then cannulated with guide cannulas aimed over the BNST and allowed to recover for a week followed by an additional week of IA alcohol exposure. All mice were then infused bilaterally with 3 μM CNO or vehicle 30-minutes prior to the start of the drinking session in a within-subjects design (**Fig. 5C**). Alcohol consumption was measured at the 24-hour timepoint. CNO infusions into the BNST significantly attenuated alcohol consumption in stressed (but not unstressed) hM3DQ mice compared to vehicle controls (**Fig. 5C**). Two-way RM ANOVA analysis of alcohol consumption revealed a significant stress x drug interaction [F_Stress x Drug_ (1, 19) = 19.25, P =0.0003]. Posthoc Sidak test revealed that CNO infusions significantly reduced alcohol consumption in stressed males but not in unstressed controls (***, P <0.0001). We observed significant reduction in water consumption in unstressed hM3DQ mice (**Fig. 5D**). Two-way RM ANOVA analysis of water consumption revealed a significant main effect of CNO [F_Drug_ (1, 19) = 5.453, P = 0.03]. Posthoc Sidak test revealed that CNO infusion significantly reduced water consumption in control but not stressed mic e (*, P<0.05). However, total fluid consumption was unaffected in either group (**Fig. 5E**). We also represented the data as difference in drinking between vehicle and CNO-injected mice for control and stress groups (**Fig. 5F**). CNO infusions decreased alcohol consumption to a greater degree in stressed male and female mice compared to unstressed controls (***, P<0.001, unpaired t-test). In summary, these results suggest that BLA^KOR^ projections to the BNST regulate social stress-escalated alcohol consumption in both male and female mice.

**Figure 5:**
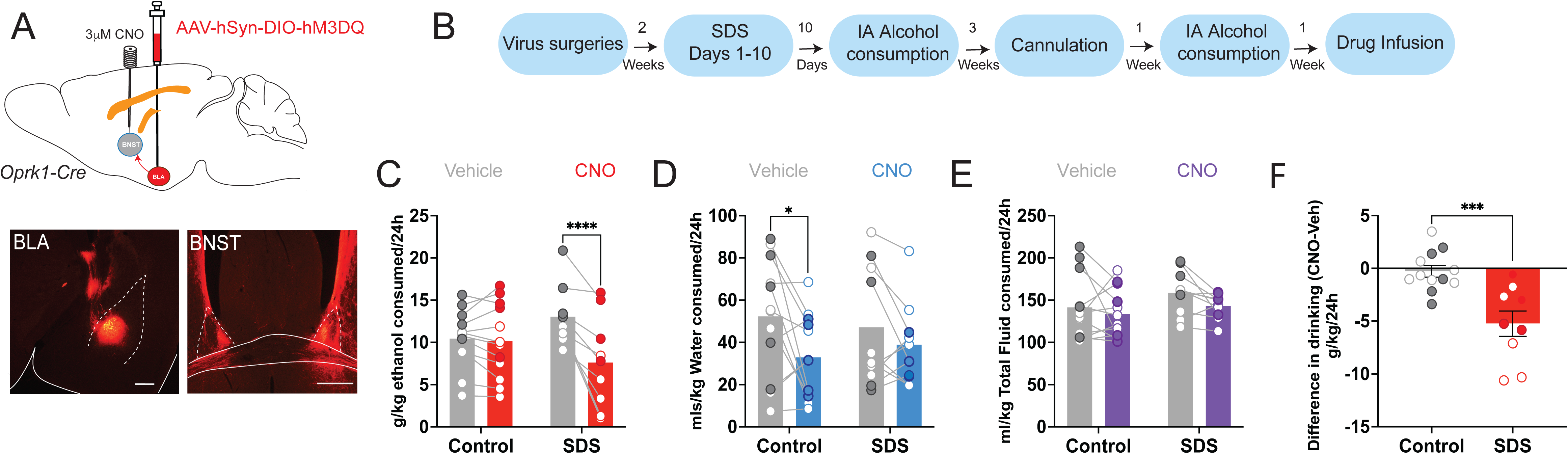
Chemogenetic activation of BLA^KOR^-BNST pathway attenuates social stress-escalated alcohol consumption in **males** and **females**. **A)** Top panel shows schematic of virus infusion and cannula placement. Bottom panels show Virus expression in BLA of *Oprk1-Cre* (bottom left) mice and BLA^KOR^ proJections to BNST (bottom right). Scalebar = 100 μm. **B)** Experimental timeline is shown. **C)** CNO infusions into the BNST significantly reduced alcohol consumption in stressed mice compared to unstressed controls. **, P<0.01, Sidak’s post-test, N = 6-7 male 4-5 female mice/group (solid color symbols). **D**) There was a significant reduction in water consumption in control hM3DQ injected mice infused with CNO. *, P<0.05, Sidak’s post-test. **E**) Total fluid consumption was not affected. **F)** Change in alcohol consumption from respective vehicle alcohol intake in control and stress grouse is shown. CNO infusions caused a greater reduction in alcohol intake in stressed mice compared to unstressed controls. ***, p<0.001, Sidak’s post-test, N= 6-7 male and 4-5 female mice/group.

## Discussion

In this study we present evidence implicating the Dyn/KOR system in regulating social stress - escalated alcohol consumption. Repeated exposure to SDS robustly and reliably increased alcohol consumption and preference in both male and female C57BL/6J mice. Stress-escalated drinking was attenuated by systemic administration of the selective KOR antagonist NorBNI in stressed male and female mice but not in unstressed controls. We used the recently generated *Oprk1-Cre* mice to determine brain regions and circuits in which KORs are functioning to regulate SDS-escalated drinking. Focusing on the BLA, we found that chemogenetic activation of BLA^KORs^ in stressed male mice significantly attenuated stress-escalated alcohol consumption thereby implicating these cells in regulating stress-escalated drinking. BLA^KOR^ neurons project robustly to the anterodorsal subdivision of the BNST, a brain region known to be important for mediating the behavioral effects of stress (44, 46, 55). We found that BNST-specific antagonism of KORs also attenuated SDS-escalated drinking. Further, chemogenetic activation of BLA^KOR^ projections to the BNST also attenuated SDS-escalated drinking in both sexes. In summary, these results suggest a role for KORs located on BLA terminals in the BNST in regulating SDS-escalated drinking.

Contrary to several published reports suggesting female mice consume greater amounts of alcohol in the IA alcohol procedure than their male counterparts (48), females in our study consumed similar amounts of alcohol compared to males. One reason for this discrepancy could be that that initial cohorts of female mice in our study were maintained on TL2019S chow which resulted in overall lower levels of baseline drinking but robust stress-escalated drinking. Later cohorts were switched to LD5053 chow which results in higher basal levels of drinking (46). Importantly, we found that SDS significantly increased alcohol consumption in both sexes regardless of the type of chow used. Hence, we combined data from these two cohorts of mice which resulted in overall lower levels of drinking in females.

The Dyn/KOR system has been previously implicated in regulating various aspects of alcohol misuse. Genetic variants of the *Oprk1* gene have been linked to Alcohol Use Disorder (AUD) (56). KOR antagonists can reduce alcohol consumption in alcohol-preferring P rats (57) and mice lacking KORs show reduced alcohol consumption. KOR agonists can promote (40) whereas antagonists can attenuate alcohol seeking behaviors (58). Dyn/KOR signaling in the extended amygdala contributes to forced swim stress induced increases in alcohol consumption in mice with a history of exposure to alcohol (59) as well as excessive operant alcohol consumption in alcohol-dependent rats (60). Dyn/KOR signaling in the extended amygdala also contributes to binge alcohol consumption (41, 61). The Dyn/KOR system is dysregulated in the BNST during alcohol withdrawal and intra-BNST antagonism of KORs attenuates increased alcohol consumption and negative affective behaviors observed during acute alcohol withdrawal (62). KORs in the VTA and medial prefrontal cortex contribute to select features of alcohol dependence (63) and alcohol-induced impairments in working memory (64).

Consistent with previously published reports (16, 17, 19), SDS robustly escalated alcohol consumption and preference in both males and females. Further, this increase in alcohol consumption was persistent and lasted several weeks post stress suggesting long-lasting neuroadaptations induced by repeated exposure to social stress. Systemic administration of NorBNI significantly attenuated stress-escalated alcohol consumption several weeks post stress suggesting persistent engagement of the Dyn/KOR system by SDS. There could be three potential mechanisms that underlie this long-lasting engagement of Dyn/KOR system post stress: 1) SDS could lead to increased KOR expression on BLA terminals 2) stress could trigger persistent Dyn release in the BNST 3) stress could lead to persistent activation of KORs located on BLA terminals in the BNST. Persistent activation of Dyn/KOR systems by stress has been previously reported. A single exposure to swim stress results in a long-lasting block of long-term potentiation at GABAergic synapses (LTP_GABA_) in the VTA (65, 66). This LTP_GABA_ block was mediated by constitutive activation of KORs located on VTA GABA neurons (65, 66). Intra-VTA antagonism of KORs several days post stress relieved this LTP_GABA_ block and attenuated stress-induced reinstatement to drug seeking (65, 66). However, this phenomenon has so far only been demonstrated at GABAergic synapses in the VTA and is absent from glutamatergic synapses. It remains to be determined if such modulation can occur on BLA^KOR^ inputs to the BNST. Adolescent social isolation stress (SI) leads to increased alcohol consumption in adults via a KOR-dependent mechanism. SI leads to long-lasting functional upregulation of the Dyn/KOR system and consequent reductions in nucleus accumbens (NAc) DA release which leads to escalation of alcohol intake in adulthood (67). NorBNI administration in adulthood reduced SI-escalated alcohol intake (67). Future studies will use electrophysiological techniques combined with optogenetics in *Oprk1-Cre* mice to determine which of these mechanisms underlie persistent engagement of Dyn/KOR system post-stress.

KORs are widely expressed in the brain in several brain areas implicated in regulating drug seeking and taking (26, 43, 45). To determine brain regions within which KORs are functioning to regulate stress-escalated drinking, we decided to make use of the newly generated *Oprk1-Cre* mouse line (46). We first examined the BLA because of its well-established role in regulating the behavioral effects of stress (49, 50), and alcohol consumption (51, 52), and high levels of KOR expression (51). We posited that stress induced increases in Dyn would inhibit BLA^KOR^ cells. To test this hypothesis, we determined the consequences of chemogenetic activation of BLA^KOR^ cells on stress-escalated alcohol consumption. We found that chemogenetic activation of BLA^KOR^ cells significantly attenuated SDS-escalated alcohol consumption in males. We also found that alcohol consumption was also significantly reduced in non-stressed control males. These results are consistent with a role for KORs in regulating basal alcohol consumption (41). A recent study found that stimulation of KORs in the NAc shell can affect alcohol drinking in a subregion and sex dependent manner (53). Hence it is possible that BLA^KOR^ inputs to the NAc shell modulate basal alcohol consumption. Our results implicate a general role for KORs in mediating alcohol reinforcement in addition to stress-escalated alcohol consumption.

BLA^KORs^ project to several brain regions including the medial prefrontal cortex (mPFC), ventral hippocampus, NAc lateral shell, and the BNST. There is evidence that BLA projections to the BNST are modulated by Dynorphin (44). Activation of KORs inhibits glutamate release from BLA inputs to the BNST and occludes the anxiolytic phenotype observed upon optogenetic activation of BLA inputs to the BNST. In summary, these results indicate that local Dyn release in the BNST can inhibit an anxiolytic pathway via activation of KORs located on BLA terminals in the BNST (44). We sought to determine whether BLA^KOR^ projections to the BNST regulate social stress-escalated drinking. BNST-specific KOR antagonism significantly attenuated social stress-escalated alcohol consumption. Further, activation of BLA^KOR^ inputs to the BNST attenuated stress-escalated alcohol consumption in both males and females. Contrary to our results with systemic CNO injections, pathway specific activation of BLA^KOR^ cells did not impact alcohol consumption in control unstressed mice suggesting that this pathway does not regulate basal alcohol consumption. These results indicate the importance of this pathway in regulating social stress-escalated alcohol intake in both sexes. One potential caveat with these results is that while it establishes a role for BLA^KOR^ projections to the BNST in stress-escalated drinking, it does not rule out a role for BLA^KOR^ projections to other brain regions including the NAc and medial prefrontal cortex (46). It is possible that BLA^KOR^ projections to multiple brain regions regulate social stress-escalated drinking. Future studies will dissect the contributions of other BLA^KOR^ inputs to stress-escalated drinking. There is evidence that KORs located on BLA terminals can modulate glutamate release. One recent study found that BLA^KOR^ inputs to the dorsomedial striatum sculpts goal directed behavior. This study found that glutamate release from BLA^KOR^ axons in the DMS evoked pDyn release in the striatum which resulted in retrograde inhibition of BLA^KOR^ terminals in the DMS (68).

KOR expression and function in the BNST is sexually dimorphic. Sex differences have been reported in the expression KOR mRNAs in the BNST. Further, administration of KOR agonist U50,488 leads to greater immediate early gene activation in females compared to males (69, 70). However, we observed no sex differences in the stress-induced engagement of KORs in the BLA-BNST pathway. SDS robustly increased alcohol consumption in both males and females which was significantly attenuated by systemic administration of the selective KOR antagonist NorBNI in both sexes. Further, chemogenetic activation of BLA^KOR^-BNST pathway attenuated stress-escalated alcohol consumption in both sexes.

Our results suggest a scenario where exposure to repeated SDS leads to Dyn release in the BNST which then acts on KORs located on BLA^KOR^ terminals in the BNST leading to inhibition of glutamate release. What is the source of Dyn in the BNST that is recruited by exposure to SDS? There are four potential sources of pDyn into the BNST: 1) CeA (59), 2) DRN (71), 3) lateral hypothalamus, and 4) local pDyn cells in the BNST (44). Future studies will determine which of these sources of pDyn into the BNST are recruited by SDS. DRN pDyn cells are particularly intriguing in the context of maladaptive social behaviors. A recent study suggested a role for this population of cells in mediating social deficits during opioid withdrawal (71). While this study showed strong projections from DRN^Dyn^ cells to the BNST, this projection has never been studies in the context of alcohol consumption.

In summary, our results have uncovered a novel role for KORs expressed on BLA glutamatergic terminals in the BNST in regulating social stress-escalated alcohol consumption. Several details regarding the source of Dyn in the BNST that is recruited by SDS and the precise temporal sequelae of Dyn release and subsequent engagement of BLA^KOR^ terminals in the BNST remain unknown and will be the focus of future studies. Our results nevertheless indicate that KOR antagonism could be a viable treatment strategy for patients suffering from comorbid social stress and AUD.

## Acknowledgements

The authors would like to thank Matthew Brian Pomrenze, Tiffany Wills, and Nicholas Gilpin for many helpful discussions on this work and for reading and providing valuable feedback on early drafts of this manuscript. This work was supported by 1R01AA027293 from NIAAA and LSUHSC startup funds.

## Notes

### Competing Interest Statement

The authors have declared no competing interest.

