## Supplementary figure legend for "BLA^KOR^ inputs to the BNST regulate social stress-escalated alcohol consumption"

**Figure S1:** The magnitude of stress-escalated drinking is similar between males and females. % Increase in drinking per 24h session in stressed mice compared to controls across 3-weeks of drinking is shown. Significant differences were not observed in the magnitude of stress effects on alcohol consumption. N = 26-35 mice/group for males and 23-24 mice/group for females.
