## Supplementary figures and images for "BLA^KOR^ inputs to the BNST regulate social stress-escalated alcohol consumption"

### Supplementary Figure 1

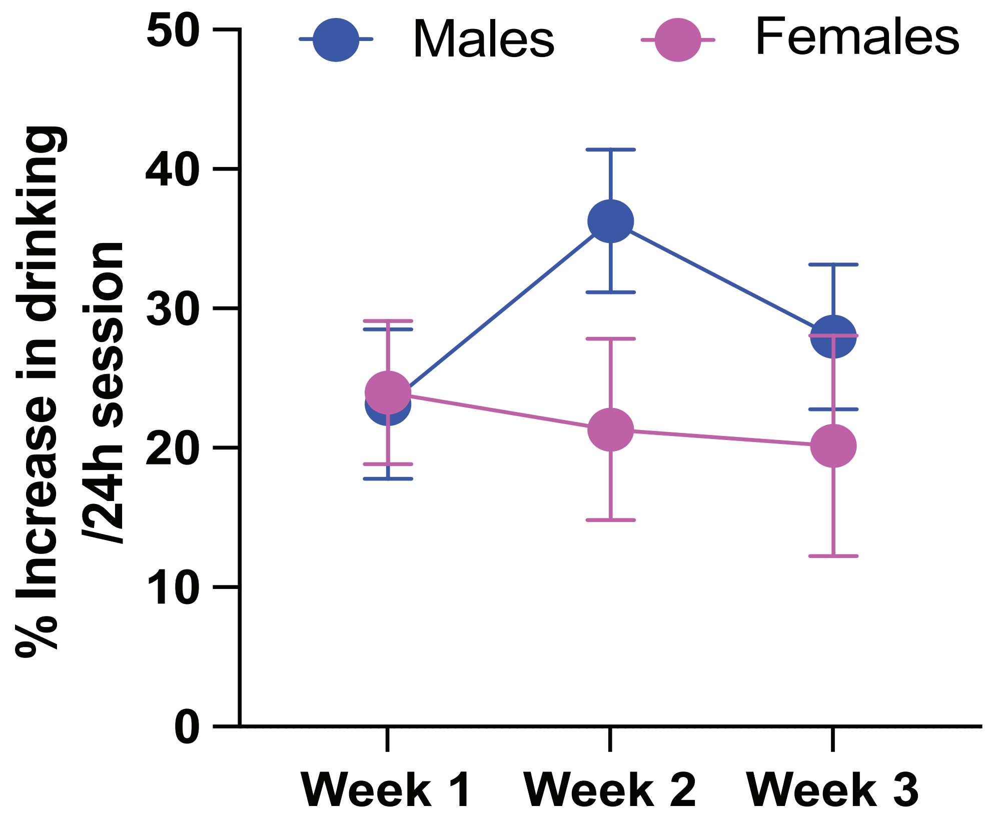
